# A wheat immune receptor pair composed of NLR and MLKL confers stable resistance to pathotypes of the blast fungus by recognizing three effectors

**DOI:** 10.64898/2026.08.07.743458

**Authors:** Soichiro Asuke, Reina Tsuchiya, Hiroyasu Kano, Fumitaka Abe, Mitsuko Kishi-Kaboshi, Mei Monta, Yuta Umehara, Mizuki Iwakawa, Harumi Koike, Yoshihiro Matsuoka, Motoki Shimizu, Yukio Tosa

## Abstract

Kinase fusion proteins (KFPs) have emerged as an important group of immune receptors encoded by plant resistance genes. Here, we report a new type of gene pair that controls resistance of wheat to the blast fungus, *Pyricularia oryzae*. We cloned a fungal gene involved in avirulence of *P. oryzae* pathotype *Eleusine* on wheat and designated it *PWT8*. We also identified its corresponding resistance gene in wheat, and tentatively named it *Rwt8*. This resistance gene was located at the same locus as previously identified resistance genes *Rwt3* and *Rwt6*. Molecular cloning revealed that *Rwt3*, *Rwt6*, and *Rwt8* were the same gene consisting of an identical gene pair, one encoding an NLR and the other encoding a mixed lineage kinase-like (MLKL) protein. These two genes were closely linked in a head-to-head orientation and behaved as a single gene. This gene pair recognized three AVR genes, *PWT3*, *PWT6*, and *PWT8*, and was designated *Rwt3.6.8*. The distribution of *Rwt3.6.8* in common wheat landraces suggested that the gene pair may have been a factor which the D genome provided to the genus *Triticum* to broaden its adaptability to various environments in the world, especially in Asia and Africa.

## Introduction

Plants carry many resistance genes (R genes) to defend themselves against pathogenic microbes. Most cloned R genes encode nucleotide-binding leucine-rich repeat (NLR) proteins^1,2^. NLRs recognize effectors encoded by corresponding avirulence genes (AVR genes) through direct or indirect binding and activate immune responses^2–5^. Some NLRs function in pairs in which one NLR (sensor) is involved in effector recognition and the other (helper) activates downstream responses leading to hypersensitive cell death^5^. Recently, kinase fusion proteins (KFPs) have emerged as another important group of immune receptors encoded by R genes^6,7^. A well-studied subgroup of KFPs is a family of tandem kinase proteins (TKPs), which possess two kinase and/or pseudokinase domains^1,8^. Recent studies indicated that *Sr62* and *Rwt4*, TKP genes for resistance to stem rust and blast, respectively, cooperated with the same helper NLR that directed execution of the resistance^9,10^.

*Pyricularia oryzae* (syn. *Magnaporthe oryzae*), the blast fungus of gramineous plants, comprises several host-specific subgroups such as the *Oryza* pathotype (MoO) pathogenic on rice (the rice blast fungus), the *Setaria* pathotype (MoS) pathogenic on foxtail millet, the *Eleusine* pathotype (MoE) pathogenic on finger millet, the *Lolium* pathotype (MoL) pathogenic on perennial ryegrass, the *Avena* pathotype pathogenic on oats, and the *Triticum* pathotype (MoT) pathogenic on wheat (the wheat blast fungus)^11–13^. MoE is further divided into phylogenetically distinct subgroups, EC-I and EC-II^14^. MoT first emerged in Brazil in 1985^15^ and subsequently spread to neighboring countries in South America^16^. It further spread to Bangladesh in 2016^17,18^, and to Zambia in 2018^19^, and is now considered a pandemic disease^20^. Inoue et al.^21^ suggested that MoT evolved from MoL or its relatives by loss of function of AVR gene *PWT3*. To further investigate the history of pathotype differentiation, we performed genetic analyses of an F_1_ population derived from a cross between an MoE (EC-II) isolate and an MoT isolate, and found that at least five genes are involved in avirulence of MoE on common wheat (*Triticum aestivum* L.)^22^. This finding implies that the loss of at least five AVR genes was required for a common ancestor of MoE and MoT to acquire virulence on common wheat. One of the five genes was identified as a variant of *PWT3*^22^. Subsequently, we isolated the second AVR gene and designated it *PWT6*^23^. Furthermore, we found that the third AVR gene was a variant of *PWT7* involved in the avirulence of an *Avena* isolate on common wheat^24^. We also identified R genes corresponding to *PWT3*, *PWT6*, and *PWT7* in common wheat, and designated them as *Rwt3*, *Rwt6* and *Rwt7*, respectively^21,25,26^. They were all located in the D genome that was provided by *Aegilops tauschii* Coss. in the evolution of common wheat^27,28^. Molecular cloning revealed that *Rwt3* and *Rwt7* encoded an NLR and a TKP, respectively^26,29^.

In the present study we cloned the fourth gene involved in the avirulence of MoE on common wheat, and designated it *PWT8*. We also identified its corresponding R gene in common wheat, and tentatively named it *Rwt8*. Interestingly, this R gene was located at the same locus as *Rwt3* and *Rwt6*. Molecular cloning revealed that *Rwt3*, *Rwt6*, and *Rwt8* were the same Mendelian unit consisting of two genes encoding NLR and KFP (other than TKP) closely linked in a head-to-head orientation and behaving as a single gene. This gene pair corresponded to three AVR genes, *PWT3*, *PWT6*, and *PWT8*. Here, we report the cloning of this new R gene pair and discuss its possible role in the geographical expansion of common wheat.

## Results

### Cloning of *PWT8*

MZ5-1-6 (MoE isolate belonging to the EC-II subgroup) is avirulent on common wheat cultivars, Chinese Spring (CS), Norin 4 (N4), and Hope whereas Br48 (MoT isolate) is virulent (Fig. 1a). The gene pairs identified so far in this system are illustrated in Extended Data Fig. 1. The five AVR genes identified in MZ5-1-6 were tentatively designated *eA1*, *eA2*, *eA3*, *eA4*, and *eA5*^22^. Their corresponding R genes were tentatively designated *R1*, *R2*, *R3*, *R4*, and *R5*, respectively. CS and N4 recognized all five AVR genes, and were assumed to carry all five R genes. On the other hand, cv. Hope recognized only two (*eA1* and *eA2*) of the five AVR genes, and was assumed to carry only two (*R1* and *R2*) of the five R genes. *eA1*, *eA3*, and *eA5* were isolated, and proved to be variants of, or designated as, *PWT7*, *PWT3*, and *PWT6*, respectively^22–24^. Their corresponding R genes, *R1*, *R3*, and *R5*, were detected through segregation analyses and designated *Rwt7*, *Rwt3*, and *Rwt6*, respectively. *Rwt7* was located on chromosome 7D, and *Rwt3* and *Rwt6* were closely linked on chromosome 1D^25,26^.

**Fig. 1.**
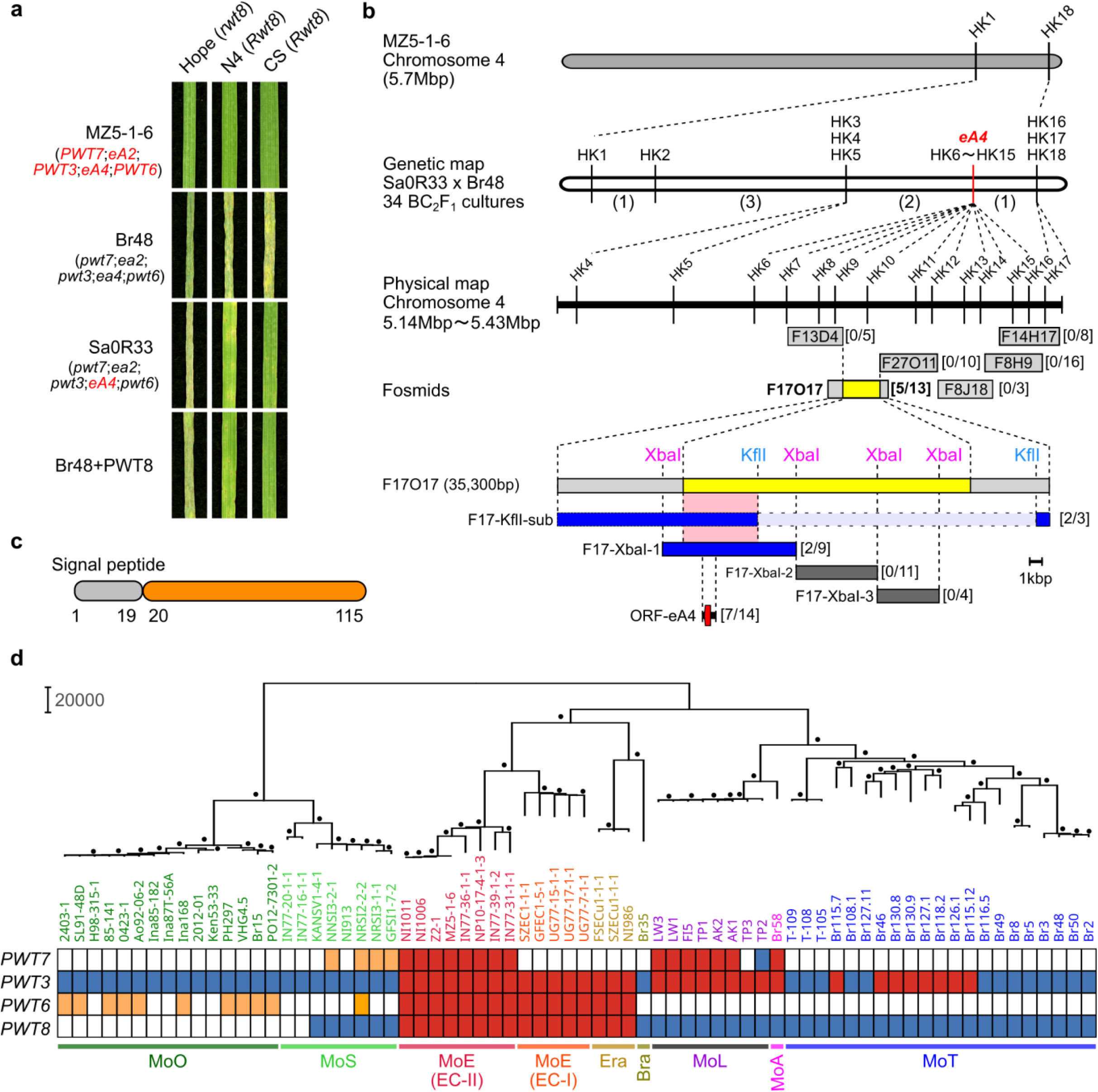
Cloning and characterization of *PWT8*. (a) Responses of wheat cultivars to MZ5-1-6 (MoE isolate), Br48 (MoT isolate), Sa0R33 (a BC₁F₁ culture), and Br48+PWT8 (a Br48 transformant carrying the *PWT8* candidate gene). (b) Genetic and physical maps around *eA4*. Numbers of recombinants are shown in parentheses. The candidate region or genes were narrowed to ∼20kb (highlighted in yellow), ∼5kb (highlighted in pink), and finally to a single gene (highlighted in red) by infection assays with transformants carrying fosmids, F17O17 subclones, and a 2kb F17-XbaI-1 subclone, respectively. Numbers in square brackets indicate the number of avirulent transformants / the total number of transformants tested. (c) Structure of the putative product of *PWT8* (*eA4*). (d) Distribution of *PWT8* and other *PWT* genes (*PWT3*, *PWT6*, and *PWT7*) in *P. oryzae*. The phylogenetic tree was constructed using the neighbor-joining method based on 490,516 biallelic SNPs. Dots indicate nodes with bootstrap support values higher than 90 (from 1,000 replicates). The tree includes 70 isolates representing MoO (*Oryza* pathotype), MoS (*Setaria* pathotype), MoE (*Eleusine* pathotypes EC-I and EC-II), Era (*Eragrostis* isolates), Bra (*Brachiaria* isolate), MoA (*Avena* pathotype), MoL (*Lolium* pathotype), and MoT (*Triticum* pathotype). Red, orange, blue, and white indicate functional type, intermediate type, non-functional type, and deletion, respectively.

To isolate *eA4*, an F_1_ culture (200R72) derived from MZ5-1-6 x Br48 was backcrossed with Br48. A BC_1_F_1_ culture (Sa0R33), which carried *eA4* alone and was avirulent on N4 and CS but virulent on Hope (Fig. 1a), was further backcrossed with Br48, resulting in a BC_2F1_ population in which *eA4* alone is segregating in an almost uniform genetic background (Extended Data Fig. 2). Infection assays on N4 revealed that avirulent and virulent cultures (*eA4* carriers and noncarriers, respectively) segregated in a 1:1 ratio (17:17) in this population. Ten cultures were arbitrarily chosen from each, bulked, and subjected to DNA sequencing. When short reads obtained were aligned to a pseudo-reference sequence of Sa0R33 generated from the genome sequence of MZ5-1-6 (GCA_004346965.1), we found a peak of the ΔSNP-index^30^ around the distal region of chromosome 4 (Extended Data Fig. 3). Among molecular markers designed in this region, ten co-segregated with the phenotype controlled by *eA4* (Fig. 1b). We screened a fosmid library of MZ5-1-6 with those markers, and constructed a contig composed of 6 fosmid clones (Fig. 1b). These clones were introduced into Br48, and resulting transformants were sprayed on N4. One clone (F17O17) conferred avirulence upon Br48 but the others including flanking clones (F13D4 and F27O11) did not (Fig. 1b), suggesting that *eA4* was located on the ∼20kb region that was present in F17O17 but absent in F13D4 and F27O11. Subcloning of F17O17 and infection assays with transformants carrying the subclones further narrowed the candidate region to 5kb (Fig. 1b). This region contained only one gene that showed a difference in expression between MZ5-1-6 and Br48 (Extended Data Fig. 4a). Its putative product was composed of 115 amino acids and had a signal peptide at its N terminus (Fig. 1c). A ∼1kb fragment containing this gene was amplified from F17O17, cloned into pBluescriptSK(+), and introduced into Br48. Infection assays revealed that transformants carrying this clone gained avirulence on N4 and CS but not on Hope (Fig. 1a, b). From these results we concluded that this gene was *eA4*, and designated it *PWT8*. Its nonfunctional allele (*pwt8*) in Br48 harbored numerous nucleotide substitutions (Extended Data Fig. 4b), and was not expressed during infection (Extended Data Fig. 4a). Functional *PWT8* was present in MoE and *Eragrostis* isolates but absent in the other pathotypes (Fig.1d).

### Identification and cloning of *Rwt3.6.8*

CS and N4 were resistant to Br48+PWT8 (a transformant of Br48 carrying the *PWT8* transgene), but Hope was susceptible (Fig. 1a). CS and N4 were crossed with Hope to identify *R4* corresponding to *PWT8*. The CS x Hope and N4 x Hope F_2_ seedling populations segregated in 3 resistant: 1 susceptible ratios whereas the N4 x CS cross yielded no susceptible F_2_ seedlings (Extended Data Table 1). These results suggest that the resistance of CS and N4 is controlled by the same, single gene. This R gene is apparently *R4* corresponding to *PWT8*, and therefore, tentatively named *Rwt8*.

A mapping population comprising 92 F_2:3_ lines from N4 x Hope was produced to identify the chromosomal location of *Rwt8*. Infection assays with Br48+PWT8 identified 19 homozygous resistant, 45 heterozygous, and 28 homozygous susceptible lines, fitting a 1:2:1 ratio. Tentative mapping with SSR markers^31^ suggested that *Rwt8* was located in the short arm of chr.1D and flanked by two SSR markers (Fig. 2a). Further mapping with additional markers narrowed the candidate region to 439 kb flanked by YU47 and RT5 (Fig. 2a). This region contained *Rwt3* encoding an NLR^29^ and a gene encoding a kinase. We tentatively designated the NLR and kinase as R3NLR (Rwt3-NLR) and R3AK (Rwt3-Associated Kinase). R3AK was a mixed lineage kinase-like (MLKL) protein^32^ with a 4-helical bundle (4HB) domain at its N terminus (Fig. 2c). *R3NLR* and *R3AK* were closely linked (2,427bp apart) in a head-to-head orientation (Fig. 2b) and segregated together in the mapping population (Extended Data Fig. 5a). In addition, they perfectly co-segregated with the phenotypes conferred by *Rwt8* (Extended Data Fig. 5a). To clarify which gene was *Rwt8*, we conducted protoplast assays based on a luciferase reporter for cell viability. Barley protoplasts were co-transfected with vectors containing a luciferase gene, *PWT8*, and *Rwt8* candidates. Luminescence was not reduced in either *PWT8-R3NLR* or *PWT8-R3AK* combinations (Fig. 2d), but when protoplasts were co-transfected with *PWT8*, *R3NLR*, and *R3AK*, luminescence was significantly reduced. These results suggest that *Rwt8* is a gene pair consisting of *R3NLR* and *R3AK*, and that both genes are required to express its function as an R gene.

**Fig. 2.**
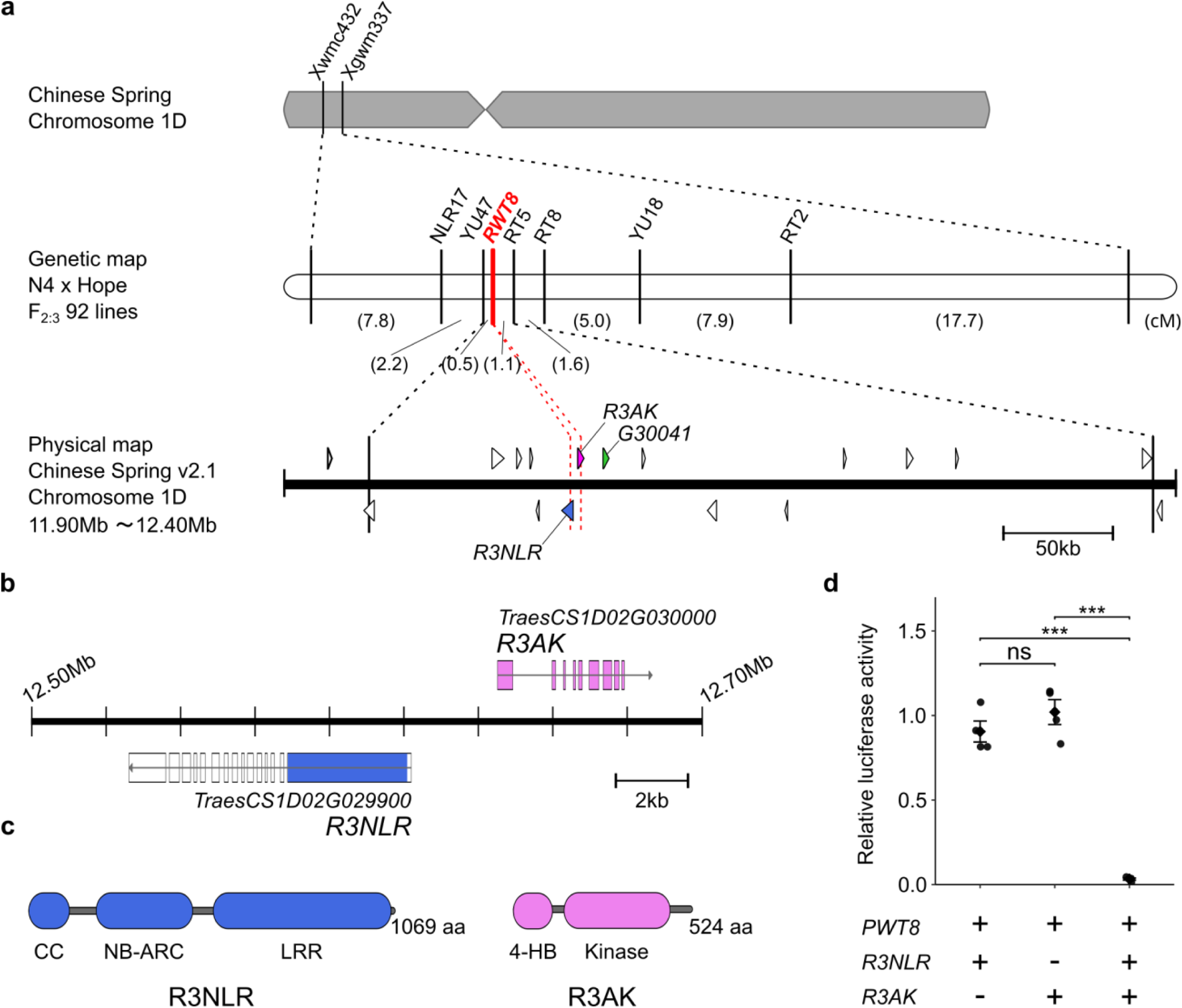
Cloning of *Rwt3.6.8.* (a) Genetic map around *RWT8* on chromosome arm 1DS constructed using N4 x Hope F_2:3_ lines. Arrowheads on the physical map indicate high-confidence genes. Genes encoding R3NLR and another NLR protein are shown in blue and green, respectively, and a gene encoding R3AK is shown in pink. (b) Gene structures of *R3NLR* (Traes1D02G029900) and *R3AK* (Traes1D02G030000) in CS. (c) Domain structures of R3NLR and R3AK. (d) Cell death assay with protoplasts. Protoplasts isolated from primary leaves of barley cv. Golden Promise were transfected with pAHC17-LUC containing a luciferase gene, pZH2Bik containing avirulence gene *PWT8* lacking the signal peptide or no insert (empty vector), and pZH2Bik vectors containing *R3NLR* cDNA, *R3AK* cDNA, or both. Luciferase activity was determined 18 h after transfection and represented as relative activities compared with those in samples with the empty vector. The experiments were repeated eight times. Results from four replicates are represented. Triple asterisks indicate significant differences at the 0.1% level. ns, not significant.

### Renaming Rwt3, Rwt6, and Rwt8 as Rwt3.6.8

*Rwt8* was mapped to a region flanked by the same marker loci (*Xwmc432* and *Xgwm337*) as *Rwt6* and *Rwt3*^25^. We hypothesized that these three are the same gene composed of *R3NLR* and *R3AK*. To test this hypothesis, the N4 x Hope mapping population (92 F_2:3_ lines) was inoculated with a Br48 transformant carrying *PWT6* (Br48+PWT6) and another transformant carrying *PWT3* (Br48+PWT3). Phenotypes conferred by Br48+PWT6 perfectly co-segregated with *R3NLR* and *R3AK* (Extended Data Fig. 5a), supporting the hypothesis that *Rwt6* is composed of the same gene pair as *Rwt8*. Phenotypes conferred by Br48+PWT3 almost co-segregated with these genes although 8 F_2:3_ lines carrying no N4 allele (of *R3NLR* and *R3AK*) showed the heterozygous phenotype (Extended Data Fig. 5a).

Transformable wheat lines lacking *Rwt3*, *Rwt6*, and *Rwt8* are needed to verify the hypothesis that the three genes are identical. Since the highly transformable common wheat cultivar Fielder (Fld) was resistant to Br48+PWT3, Br48+PWT6, and Br48+PWT8, we crossed it with cv. Transfed (Tfed) susceptible to all three, backcrossed the F_1_ with Fld, and self-fertilized a BC_1_F_1_ plant (Extended Data Fig. 6a). From the resulting BC_1_F_2_ population, FTF2723, which was highly susceptible to all three fungal transformants, was chosen for wheat transformation. cDNA of *R3NLR* and *R3AK* amplified from CS were introduced into pZH2Bik, and established as pZH-R3NLR and pZH-R3AK, respectively. They were introduced into FTF2723 separately through *Agrobacterium*-mediated transformation, and a T1 plant carrying heterozygous pZH-R3NLR was crossed with a T_1_ plant carrying homozygous pZH-R3AK (Extended Data Fig. 6b). When selected homozygous F_2_ transformants carrying both clones, pZH-R3NLR alone, pZH-R3AK alone, and none were inoculated with Br48+PWT3, Br48+PWT6, and Br48+PWT8, those carrying pZH-R3NLR alone or pZH-R3AK alone were susceptible to all of the three fungal transformants (Fig. 3). On the other hand, the transformants carrying both clones were resistant to all of them (Fig. 3). From these results, we concluded that *Rwt3*, *Rwt6*, and *Rwt8* are the same gene composed of *R3NLR* and *R3AK*.

**Fig. 3.**
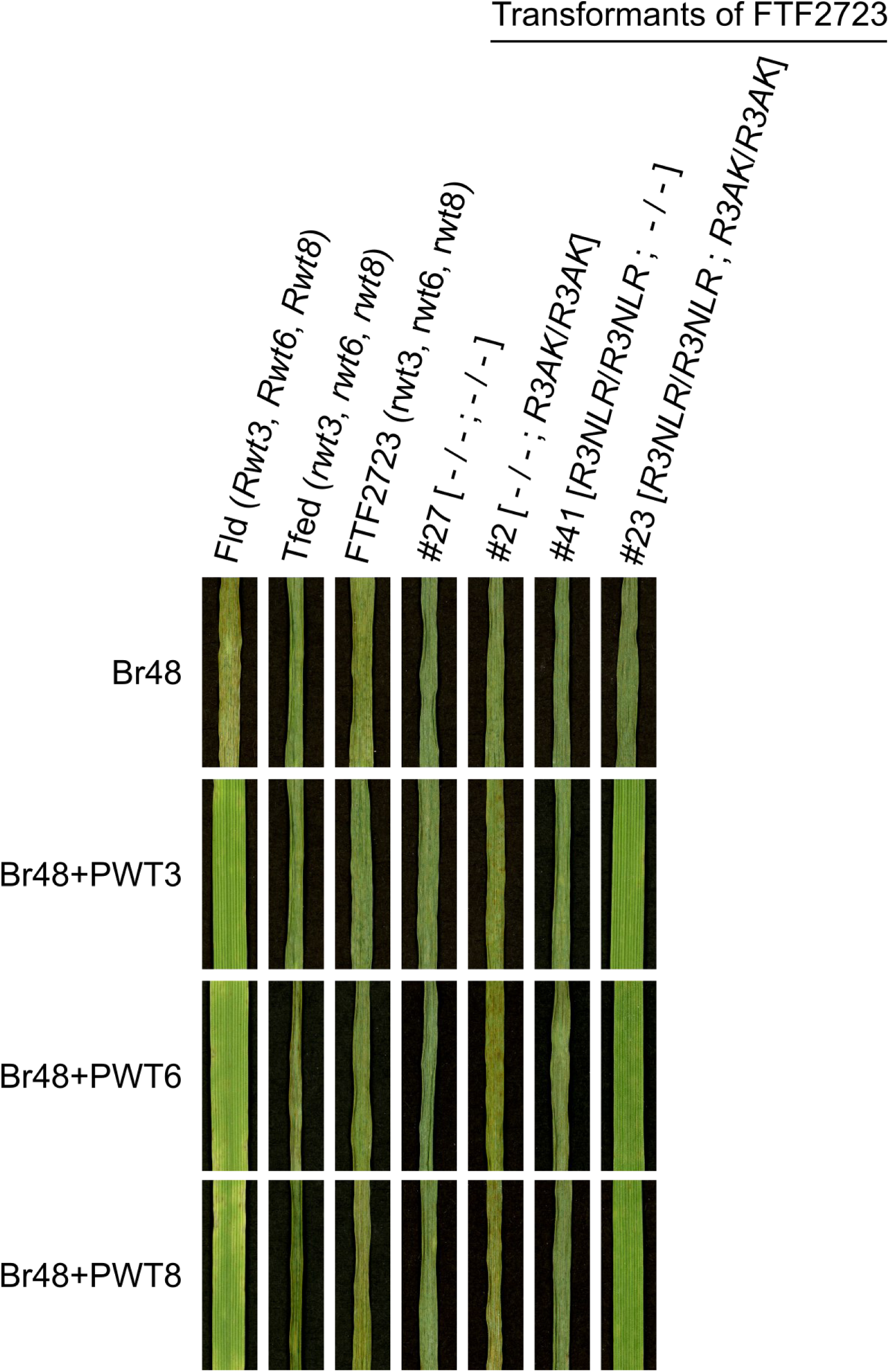
Responses of wheat transformants expressing *R3NLR* and/or *R3AK* to Br48 and its transformants carrying *PWT3*, *PWT6*, or *PWT8*. Genotypes of the parental cultivars are shown in parentheses. Transgenes carried by wheat transformants are shown in square brackets.

To clarify why several F_2:3_ lines carrying no *R3NLR* and *R3AK* N4 alleles showed the heterozygous phenotype against Br48+PWT3 (Extended Data Fig. 5a), ∼16 seedlings of some of these F_2:3_ lines were again inoculated with Br48+PWT3. In all cases, resistant and susceptible seedlings segregated in 3:1 ratios (Extended Data Fig. 5b). This result suggests that N4 carries an additional R gene that recognizes *PWT3* but does not recognize *PWT6* or *PWT8*.

### Genome structures around the NLR-MLKL gene pair and its geographical distribution

Genome structures around the NLR-MLKL gene pair were examined using whole-genome sequence data of representative cultivars analyzed in the 10+ genome sequence project^33^. Cultivar Norin 61 with almost the same structure as CS carried both *R3NLR* and *R3AK*, and responded to all of *PWT3*, *PWT6*, and *PWT8* (Fig. 4a). On the other hand, cv. Paragon, which did not respond to any of the three avirulence genes, had several deletions in the region and had lost *R3NLR* and *R3AK*.

**Fig. 4.**
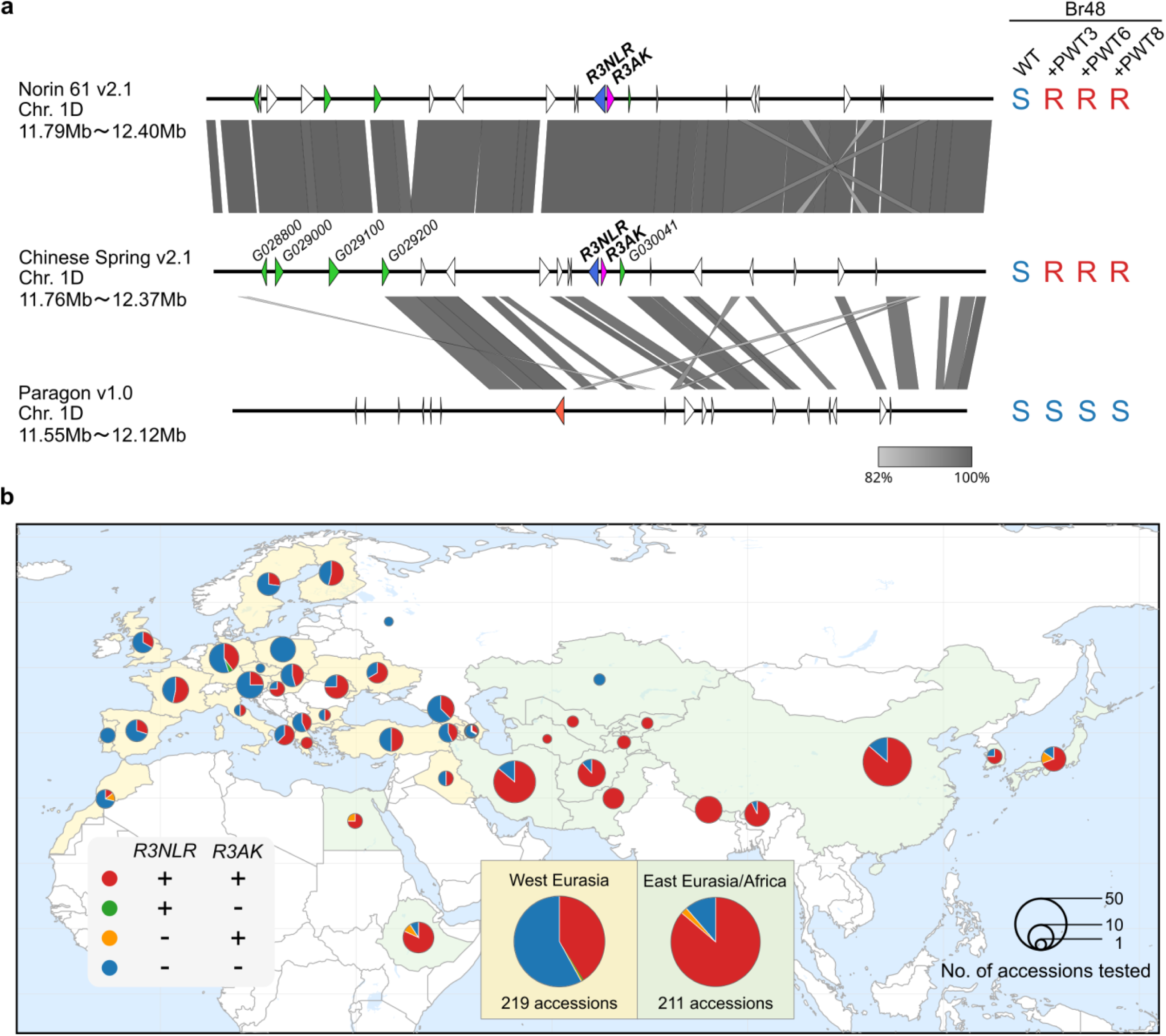
Structure and distribution of *Rwt3.6.8* in cultivars and landraces of common wheat. (a) Alignments of the genomic region around *Rwt3.6.8* in cv. Norin 61, Chinese Spring, and Paragon analyzed in the 10+ genome project. High-confidence genes in each genome are indicated by arrowheads. Genes encoding R3NLR and other NLR proteins are shown in blue and green, respectively; genes encoding R3AK and other kinase fusion proteins are shown in pink and orange, respectively. Reactions of these cultivars to Br48 and its transformants are shown at the right (R, resistant; S, susceptible). (b) Global distribution of *Rwt3.6.8* in common wheat landraces. The presence or absence of *R3NLR* and *R3AK* was determined using PCR markers R3NLR-PA and R3AK-PA, respectively. Eurasia was divided into western and eastern regions using the Caspian Sea area, where common wheat originated, as the boundary. The countries classified into western Eurasia are painted in light yellow while those classified into eastern Eurasia and Africa are painted in light green.

The geographical distribution of *R3NLR* and *R3AK* was surveyed using 431 landraces of common wheat collected worldwide. These two genes were distributed almost always together (Fig. 4b), supporting the idea that they inherit as a single Mendelian factor. In Eurasia, their frequencies showed a sharp contrast between western and eastern regions of Caspian Sea area; they were ∼40% in the western region including Europe but exceeded 80% in the eastern region (Fig. 4b). Their frequencies were also high in Africa (Ethiopia and Egypt).

## Discussion

In the present study we found a gene pair corresponding to three avirulence genes, *PWT3*, *PWT6*, and *PWT8*, and designated it *Rwt3.6.8*. This gene pair consisted of a gene encoding an NLR and a gene encoding an MLKL protein. This is a novel type of gene pair involved in resistance to pathogens. In the *Sr62* and *Rwt4* systems these TKP genes were sensors whereas the associated NLR was a helper^9,10^. Conversely, the NLR in our system is a putative sensor and the MLKL protein is a helper, which will be reported in a co-submitted paper by Bennett et al.^34^. The NLR was previously reported to be *Rwt3*^29^. From the viewpoint of Mendelian genetics, however, *Rwt3* should be the gene pair consisting of the NLR and MLKL genes because they are inherited together and behave as a single gene (Extended Data Fig. 5a, Fig.4b). Therefore, we rename the NLR gene as *R3NLR*. Using this revised nomenclature, we propose a new definition of these gene names; *Rwt3*=*Rwt6*=*Rwt8*=*Rwt3.6.8* consisting of *R3NLR* and *R3AK*. Although Asuke et al. ^25^ recognized some “recombinants” between *Rwt3* and *Rwt6* and some landraces that recognized *PWT3* but not *PWT6*, these apparent discrepancies could be attributed to another gene that was linked to *Rwt3.6.8* but recognized *PWT3* alone (Extended Data Fig. 5a, b). *Rwt3* and *Rwt6* are common names of *Rmg6* and *Rmg9*, respectively^21,25^. We propose that *Rmg9* be adopted as a formal name of *Rwt3.6.8* because the line possessing *Rmg6* was later shown to include a gene recognizing *PWT3* alone in addition to *Rwt3.6.8* as mentioned above.

We isolated the fourth avirulence gene, *PWT8*, involved in avirulence of MoE (EC-II) on common wheat. The distribution of the four genes (*PWT7*, *PWT3*, *PWT6*, and *PWT8*) in *P. oryzae* is summarized in Fig. 5. *P. oryzae* is roughly divided into two groups, one including MoO and MoS and the other including MoE, MoL, MoA, MoT, and isolates from *Eragrostis* spp.^24^. Avirulence of MoS and MoO on common wheat is mainly controlled by avirulence gene *PWT1* ^35,36^ and loss of function of the *ACE1* cluster^37^. On the other hand, the avirulence/virulence of the other group on common wheat can be explained by the four avirulence genes. *PWT3* is carried by all of the pathotypes/isolates in this group except MoT, suggesting that common wheat acquired the resistance to this group mainly through gain of *Rwt3.6.8*. If the avirulence gene involved in the resistance had been only *PWT3*, the resistance would have been easily overcome by its mutation (loss of function). However, *Rwt3.6.8* recognizes *PWT6* and *PWT8* in addition to *PWT3*. It is highly unlikely that mutations leading to the loss of function occur in all three genes simultaneously. Therefore, the gain of *Rwt3.6.8* should have conferred robust and stable resistance on wheat against MoE and *Eragrostis* isolates which carry all three genes (Fig. 5).

**Fig. 5.**
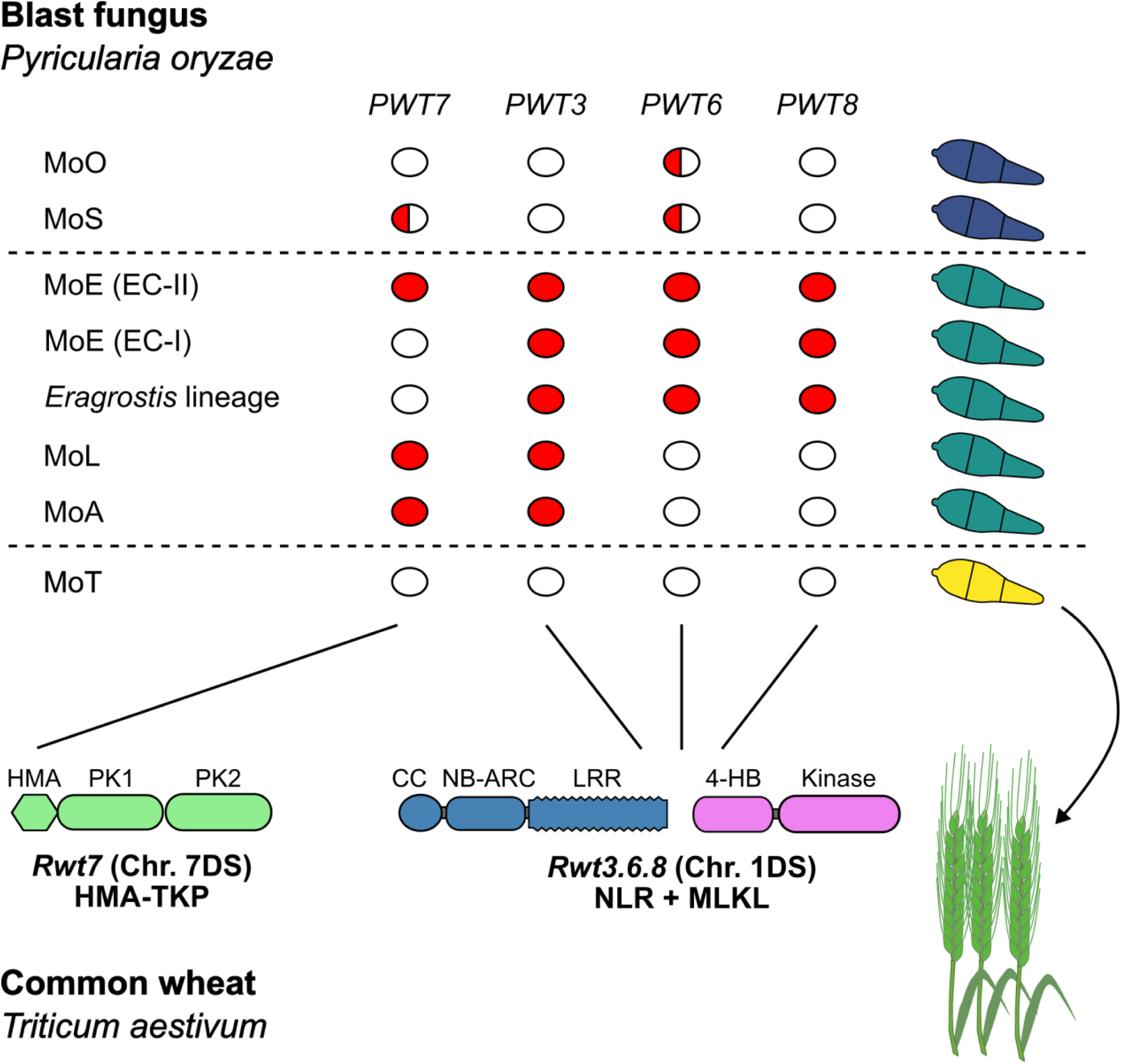
Kinase fusion proteins determine the incompatibility between non-adapted pathotypes of *Pyricularia oryzae* and common wheat (*Triticum aestivum*). Solid and half-filled red ellipses indicate ubiquitous and partial distribution, respectively, of functional *PWT* genes acting as AVR genes. MoE including EC-I and EC-II, the *Eragrostis* lineage, MoL, and MoA generally carry two or more *PWT* genes that are recognized by either *Rwt7* or *Rwt3.6.8*, resulting in avirulence. Wheat-infecting isolates (MoT) have undergone loss-of-function mutations in all *PWT* genes, thereby escaping recognition by these immune receptors.

Common wheat (AABBDD) arose through the hybridization between tetraploid wheat (*T. turgidum* L., AABB) and *Ae. tauschii* (DD)^27,28^ in a northeastern portion of the Fertile Crescent and its neighboring regions of Transcaucasus and the southern coastal Caspian^38^. The D genome conferred broader adaptability on the genus *Triticum*, allowing common wheat to spread worldwide^39,40^. It should be noted that both *Rwt7* and *Rwt3.6.8* are located on the D genome. In other words, the ability of common wheat to recognize *PWT7*, *PWT3*, *PWT6*, and *PWT8* is conferred by the D genome. We suggest that this may be an important factor which the D genome provided to the genus *Triticum* to broaden its adaptability, especially in Asia and Africa, and that *Rwt3.6.8* was a key underlying gene. This idea is supported by the distribution of *Rwt3.6.8*. Fig. 4b shows that the frequency of *Rwt3.6.8* carriers is higher in Africa and the eastern Eurasia (east of the Caspian region in which common wheat originated) than in the western Eurasia including Europe. Finger millet was domesticated in western Uganda and the Ethiopian highlands^41,42^, then spread east, and is now widely cultivated in Africa, and South and East Asia^43^. Blast is the most important disease affecting the growth and yield of finger millet^44^, implying that its causal agent, MoE, is also widely distributed in these areas. *Rwt3.6.8* may have played an important role in the survival of common wheat in these areas where finger millet and its pathogen (MoE) are widely distributed.

## Methods

### Fungal materials

*Pyricularia oryzae* isolates used in the present study are listed in Supplementary Table S1. An F₁ culture 200R72 was derived from a cross between the MoE isolate MZ5-1-6 and the MoT isolate Br48^22^. 200R72 was subsequently backcrossed with Br48 to generate BC₁F₁ cultures, from which Sa0R33 was selected as a carrier of *eA4*. Sa0R33 was further backcrossed with Br48 on oatmeal agar medium, as described by Murakami et al.^35^, and 34 BC₂F₁ cultures were randomly isolated for mapping *eA4* (*PWT8*). The mating types and genotypes at molecular marker loci of these cultures were determined by PCR using Quick Taq (TOYOBO, Japan). The primer pairs used for genotyping are listed in Supplementary Table S2. Br48 and its transformants carrying *PWT3* (strain M-16^21^), *PWT6* (strain t15^23^), or *PWT8* (strain t1, generated in the present study) were used for genetic mapping of *Rwt3*, *Rwt6*, and *Rwt8*.

### Plant materials

A total of 431 common wheat (*Triticum aestivum*) landraces provided by K. Kato, Okayama University, were surveyed to clarify worldwide distribution of *Rwt3*.6.8 using PCR-based markers, R3NLR-PA and R3AK-PA. Genomic DNA was extracted from each accession using the cetyltrimethylammonium bromide (CTAB) method. *T. aestivum* cv. Chinese Spring (CS) and Norin 4 (N4), which carry *Rwt3*, *Rwt6*, and *Rwt8*, were used as resistant controls, whereas cv. Hope, which lacks all three genes, was used as a susceptible control.

### Pathogenicity assay

Wheat seeds were sown in vermiculite supplemented with a 1:1500 dilution of liquid Hyponex fertilizer (Hyponex, Japan) in seeding containers (5.5 × 15 × 10 cm) and grown at 22°C under a 12-h light/12-h dark photoperiod for 8 days. Immediately before inoculation, the primary leaves were fixed to a plastic board with rubber bands. Conidial suspensions (1 × 10^5^ conidia/mL) containing 0.01% Tween 20 were prepared as described by Tagle et al.^45^ and sprayed onto the primary leaves using an air compressor. The inoculated seedlings were incubated in the dark under high-humidity conditions for 24 h at 22°C and then transferred to dry conditions at 22°C under a 12-h light/12-h dark photoperiod. Symptoms were evaluated 4–6 days after inoculation based on lesion size and color. Lesion size was rated on a six-point scale from 0 to 5: 0, no visible lesions; 1, pinhead-sized spots; 2, small lesions (<1.5 mm); 3, scattered, intermediate-sized lesions (<3 mm); 4, large typical lesions; and 5, complete shriveling of leaf blades. Infection types were expressed as a number indicating lesion size followed by a letter indicating lesion color: “B” for brown and “G” for green. Infection types 0 and 1B–3BG were classified as resistant, whereas infection types 3G–5G were classified as susceptible. Seedling assays were performed three times independently.

### Mapping of *eA4* by QTL-seq analysis

For bulked segregant analysis, 10 avirulent and 10 virulent cultures were selected from the 34 BC₂F₁ cultures based on their reactions on CS. Genomic DNA was extracted using the DNeasy Plant Mini Kit (QIAGEN, Germany) according to the manufacturer’s instructions. A sequencing library was prepared from each DNA sample and subjected to paired-end sequencing on an Illumina MiSeq platform to generate 2 × 300bp paired-end reads. The reads obtained from the 10 avirulent BC_2_F_1_ cultures and those from the 10 virulent BC_2_F_1_ cultures were pooled separately *in silico* to generate the *eA4* and *ea4* bulks, respectively. The pooled reads were aligned to a pseudo-reference sequence generated by substituting Sa0R33 SNPs into MZ5-1-6 reference genome. At SNP sites polymorphic between the pseudo-reference sequence and Br48, the SNP-index defined as the proportion of reads carrying the Br48 allele was calculated for each bulk using the QTL-seq pipeline^30,46^. The ΔSNP-index was calculated by subtracting the SNP-index of the *eA4* bulk from that of the *ea4* bulk. Genomic regions showing ΔSNP-index values close to the theoretical maximum (+1) were considered candidate regions linked to *eA4*.

### Construction of fosmid library of MZ5-1-6 for identifying *eA4*

To further delimit the candidate region for *eA4*, an ordered fosmid library of MZ5-1-6 was constructed using the CopyControl Fosmid Library Production Kit (Lucigen, USA) according to the manufacturer’s instructions. The library was screened by plate PCR using molecular markers targeting the *eA4* candidate region, and six positive fosmid clones (F8H9, F8J18, F13D4, F14H17, F17O17, and F27O11) were selected. Both ends of the fosmid inserts were sequenced using vector-specific primers (FP and RP) designed from the CopyControl pCC2FOS Fosmid Vector sequence (Lucigen, USA) (Supplementary Table S2). Based on the resulting end sequences, the positive clones were ordered to construct a physical contig covering the candidate region.

### Transformation of *P. oryzae*

Fosmids or pBluescript II SK(+) vectors carrying restriction enzyme-digested fosmid fragments were introduced into Br48 protoplasts by co-transformation with pSH75, which carries the hygromycin B phosphotransferase gene, as described by Tosa et al.^47^. The presence or absence of the transgene in the resulting transformants was determined by PCR analysis (Supplementary Table S2).

### RNA-seq analysis of candidate genes for *eA4*

To assess the transcription of candidate genes for *eA4* (*PWT8*), RNA-seq analysis was performed using leaves of the susceptible barley cultivar Nigrate inoculated with MZ5-1-6 (*eA4*) or Br48 (*ea4*). Leaves were collected at 24 h post-inoculation, and total RNA was extracted using the Maxwell RSC Plant RNA Kit (Promega, USA) according to the manufacturer’s instructions. Strand-specific RNA-seq libraries were prepared using the NEBNext Ultra II Directional RNA Library Prep Kit for Illumina (New England Biolabs, USA) and sequenced on an Illumina NextSeq 500 platform (Illumina, USA) to generate 2 × 150-bp paired-end reads. Adapter sequences and low-quality bases were removed using Trimmomatic v0.39^48^. The trimmed reads were mapped to the MZ5-1-6 reference genome using HISAT2 v2.1.0^49^. The resulting alignment files were converted to BAM format, sorted, and indexed using SAMtools v1.19^50^. Read coverage across the candidate loci was visualized using the Integrative Genomics Viewer (IGV)^51^.

### Distribution analysis of *PWT8* in *P. oryzae*

Whole-genome short-read data from 70 isolates listed in Supplementary Table S1 were used for phylogenetic analysis. These comprised 15 *Oryza*, eight *Setaria*, three *Eragrostis*, five EC-I *Eleusine*, eight EC-II *Eleusine*, eight *Lolium*, one *Brachiaria*, one *Avena*, and 21 *Triticum* isolates. The short reads were mapped to the MZ5-1-6 genome assembly^52^ using BWA v0.7.17^53^, and the resulting alignments were sorted using SAMtools v1.19^50^. SNPs and indels were called using the Genome Analysis Toolkit v4.1.4.1^54^, as described by Asuke et al.^23^. A neighbor-joining phylogenetic tree was constructed from the resulting biallelic SNP dataset using MEGA X, and branch support was evaluated using 1,000 bootstrap replicates^55^.

The trimmed reads were *de novo* assembled using SPAdes v4.1.0^56^. The presence and allelic variation of *PWT7*, *PWT3*, *PWT6*, and *PWT8* in each genome assembly were examined using BLASTN implemented in BLAST+ v2.6.0^57^. Sequence similarities among the corresponding genomic regions were calculated using BLASTN and visualized using Easyfig v2.2.2^58^. Genes and transposable elements within these regions were annotated by BLASTN searches using annotated genes from isolate 70-15^59^ and known transposable-element sequences as queries.

### Genetic mapping of *Rwt8*

To map *Rwt8*, 92 F₂ plants derived from a cross between N4 and Hope were grown in the field and self-pollinated to generate F_2:3_ lines. For phenotypic evaluation, 20 seeds from each F_2:3_ line were subjected to seedling inoculation assays with Br48 and Br48 transformants carrying *PWT6* (strain t15), *PWT3* (strain M-16), and *PWT8* (strain t1). Another 20 seeds from each family were grown at 22°C for 7 days, and the resulting seedlings were pooled for genomic DNA extraction using the cetyltrimethylammonium bromide (CTAB) method. Simple sequence repeat (SSR) markers reported by Somers et al.^31^ were initially screened for polymorphisms between the parental cultivars and subsequently genotyped in the F_2:3_ population. PCR amplification was performed using 2× Quick Taq HS DyeMix (TOYOBO, Japan). To develop additional markers within the candidate region, genes predicted in the Chinese Spring reference genome resources (versions 1.0, 1.1, and 2.1) were examined, with particular attention to disease resistance-related genes, including those encoding nucleotide-binding leucine-rich repeat receptors (NLRs) and protein kinases. Candidate SNPs were selected based on publicly available SNP information in Ensembl Plants (https://plants.ensembl.org/) and GrainGenes (https://graingenes.org/GG3/). Primer pairs were designed to amplify a single product from both N4 and Hope. The resulting amplicons were subjected to Sanger sequencing to verify polymorphisms between the two cultivars, and suitable polymorphisms were used to develop cleaved amplified polymorphic sequence (CAPS) markers (Supplementary Table S2). A genetic linkage map was constructed using MAPMAKER/EXP version 3.0^60^. The logarithm-of-odds (LOD) score threshold was set at 4.0, and genetic distances were calculated using the Kosambi mapping function.

### Cell death assay using barley protoplasts

Protoplast cell-death assays were performed to determine whether *R3NLR* (Traes1D02G029900) and *R3AK* (Traes1D02G030000), individually or in combination, mediate recognition of *PWT8*. Total RNA was extracted from primary leaves of CS using Sepasol-RNA I Super G (Nacalai Tesque, Japan) and treated with DNase I at 37°C for 20 min. First-strand cDNA was synthesized using the PrimeScript 1st Strand cDNA Synthesis Kit (Takara Bio, Japan). The coding sequences of *R3NLR* and *R3AK* were amplified from CS cDNA. The coding sequence of *PWT8*, excluding the region encoding the signal peptide, was amplified from MZ5-1-6 genomic DNA. The primers used for PCR amplification are listed in Supplementary Table S2. The resulting fragments were cloned into KpnI-linearized pZH2Bik using the In-Fusion Cloning Kit (Takara Bio, Japan) for expression under the control of the rice ubiquitin promoter, generating pZH-R3NLR, pZH-R3AK, and pZH-PWT8. These plasmids were purified using the NucleoBond Xtra Maxi Kit (Macherey-Nagel, Germany). Mesophyll protoplasts were prepared from the primary leaves of 8-day-old barley cv. Golden Promise (GP). Protoplast transfection was performed as described by Saur et al.^61^. Briefly, expression plasmids carrying *R3NLR* and/or *R3AK*, the *PWT8* expression plasmid, and the luciferase reporter plasmid pAHC17-LUC were mixed in equimolar amounts and co-transfected into GP protoplasts by the polyethylene glycol treatment. After incubation for 18 h at 20°C in the dark, the protoplasts were lysed. Luciferase activity in the resulting cell extracts was measured immediately after addition of the luciferase substrate using a TriStar 3 multimode microplate reader (Berthold, Germany) in luminescence mode, with an integration time of 1 s per well. Luminescence values were normalized to those of the corresponding negative control, in which the *R3NLR* and/or *R3AK* expression plasmids were replaced with equimolar amounts of the empty pZH2Bik vector. The assay was performed in four independent experiments.

### Production of transgenic plants

pZH-R3NLR and pZH-R3AK were introduced into *T. aestivum* cv. Fielder via the *Agrobacterium*-mediated transformation as described by Ishida et al.^62^. Insertions of transgenes were checked by PCR with the HPT primers (Supplementary Table 2). We obtained 10 and 7 T_1_ lines carrying *R3NLR* and *R3AK*, respectively. Transgenic T_1_ seedlings were inoculated with Br48, and its *PWT* transformants to evaluate functions of the transgenes. A T_1_ plant heterozygous for the *R3NLR* transgene (FTF2723+R3NLR #1-2-2) was crossed with a T_1_ plant homozygous for the *R3AK* transgene (FTF2723+R3AK #3-2-3) to produce F_1_ seeds. Among the resulting F_1_ plants, individuals heterozygous for both transgenes, which were expected to occur at a frequency of 1/2, were selected and self-pollinated to generate an F_2_ population. F_2_ plants were further self-pollinated to produce F_3_ lines. Using molecular markers, the following four genotypic classes were selected: (i) lines homozygous for both transgenes; (ii) lines homozygous for the R3NLR transgene and lacking the R3AK transgene; (iii) lines homozygous for the R3AK transgene and lacking the R3NLR transgene; and (iv) lines lacking both transgenes (Extended Data Fig. 6).

## Data availability

The nucleotide sequence of *PWT8* has been deposited in the DDBJ/EMBL/GenBank databases under the accession number LC942388. Raw RNA sequencing read data of *P. oryzae* are available under the accession numbers PRJDB42869 and PRJDB16420. All plasmids, plant lines, and fungal strains generated in this work are available from the authors upon request.

## Acknowledgments

We thank M. J. Banfield, John Innes Centre, UK, for useful information on molecular characteristics of the genes we isolated, Kenji Kato, Okayama University, for providing the common wheat landraces, R. A. McIntosh, University of Sydney, Australia, for valuable suggestions for gene designation and the manuscript, and Reiko Kiuchi, Kobe, for financial support. Computations were partially performed on the NIG supercomputer owned by National Institute of Genetics, Research Organization of Information and Systems. This work was supported by Grants-in-Aid for Scientific Research from the Japan Society for the Promotion of Science, 21H04726 (YT), 22K20580 (SA), and Kobe University Strategic International Collaborative Research Grant (Type B Fostering Joint Research) (SA).

## Author contributions

Y.T. and S.A. conceived the project. S.A., R.T., H. Kano, F.A., M.K., M.M., Y.U., M.I., H. Koike, and M.S. performed the experiments. Y.T, S.A., and Y.M. analyzed the data. Y.T. and S.A. wrote and revised the manuscript.

---

**Extended Data Fig. 1.**
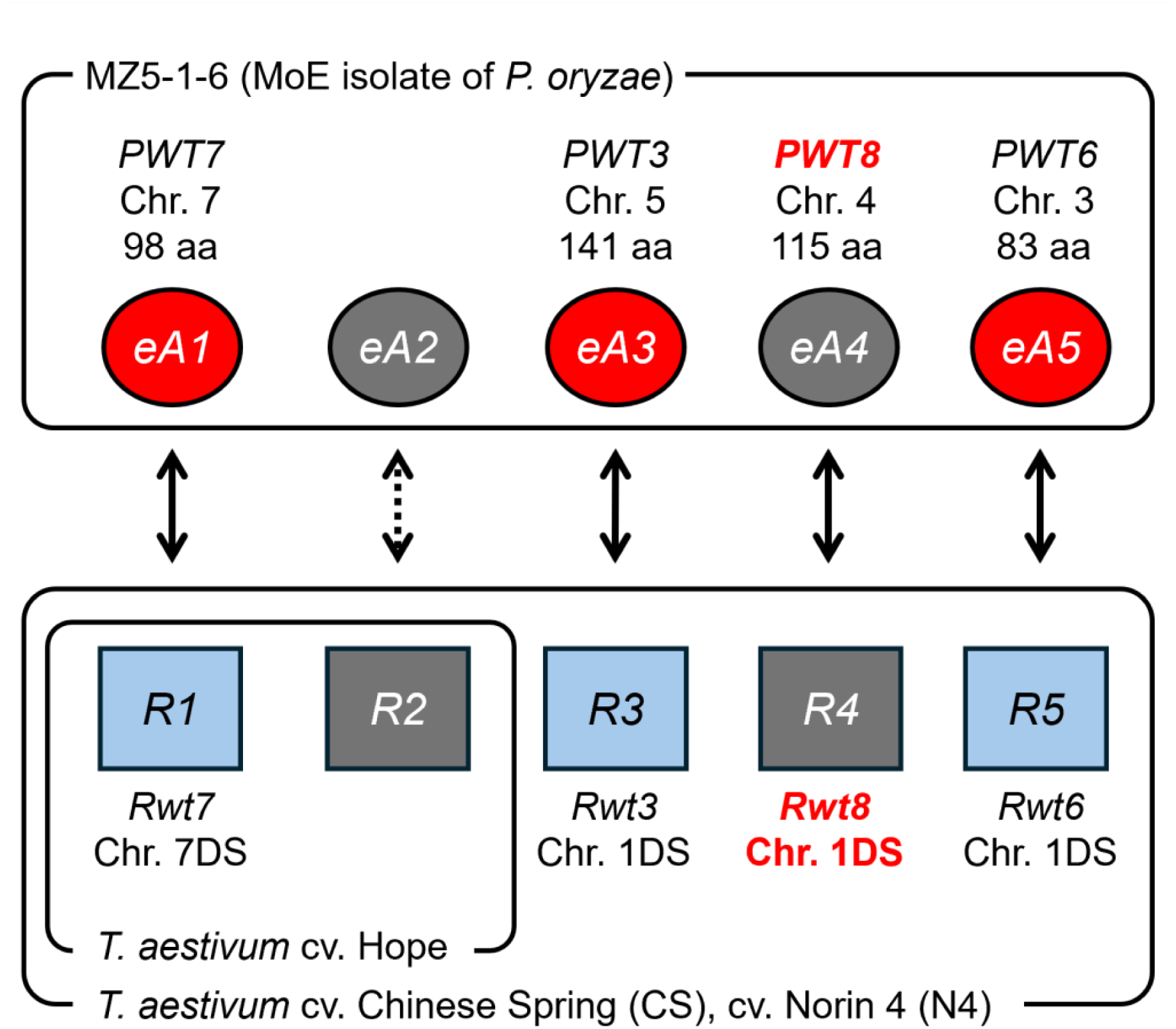
Gene-for-gene interactions between MoE (EC-II) and common wheat. Red oval - blue rectangle pairs indicate AVR gene - R gene pairs identified so far. *eA4* and *R4* were identified in the present study and designated *PWT8* and *Rwt8*, respectively.

**Extended Data Fig. 2.**
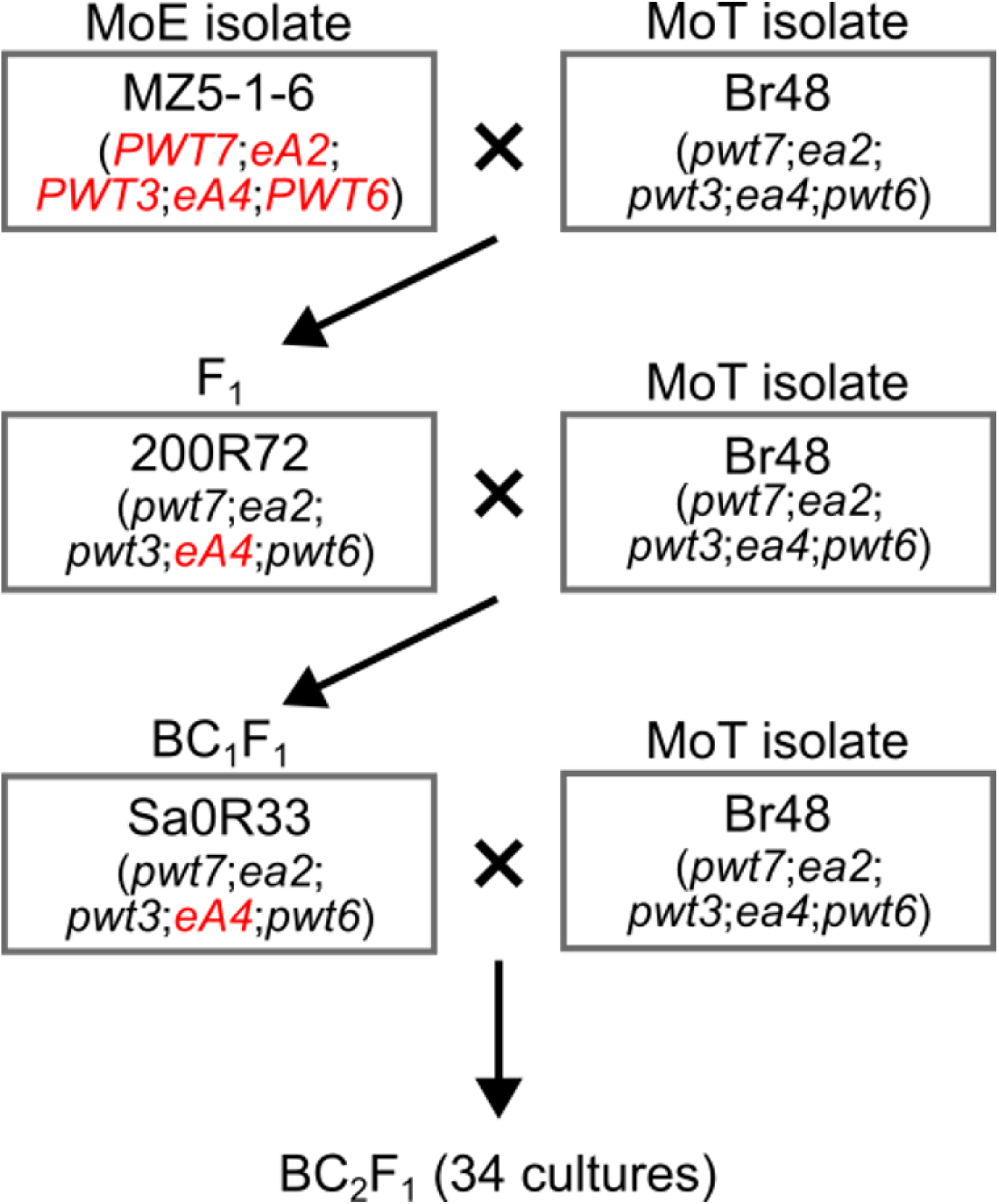
Pedigree of *P. oryzae* cultures used for cloning *eA4* (*PWT8*).

**Extended Data Fig. 3.**
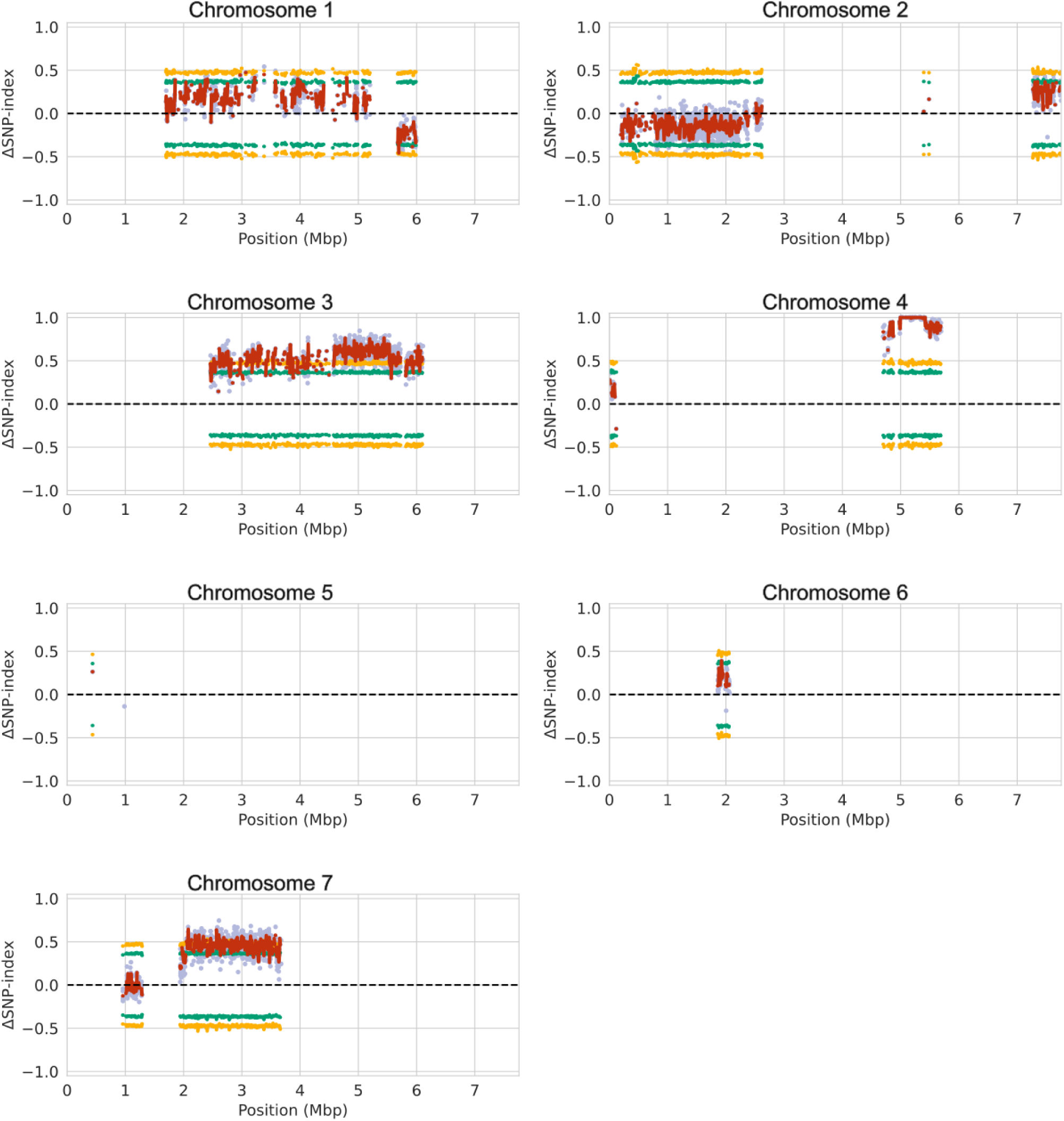
Genome-wide plots of the ΔSNP-index of avirulent and virulent bulks in the BC2F1 generation derived from Sa0R33 × Br48. Each of the avirulent (*eA4* carriers) and virulent (*ea4* carriers) bulks consisted of 10 cultures selected from the BC_2_F_1_ generation. SNP-index values in each bulk were calculated using a pseudo-reference sequence generated by substituting Sa0R33 SNPs into MZ5-1-6 reference genome. The ΔSNP-index was calculated by subtracting the SNP-index of the avirulent bulk from that of the virulent bulk.

**Extended Data Fig. 4.**
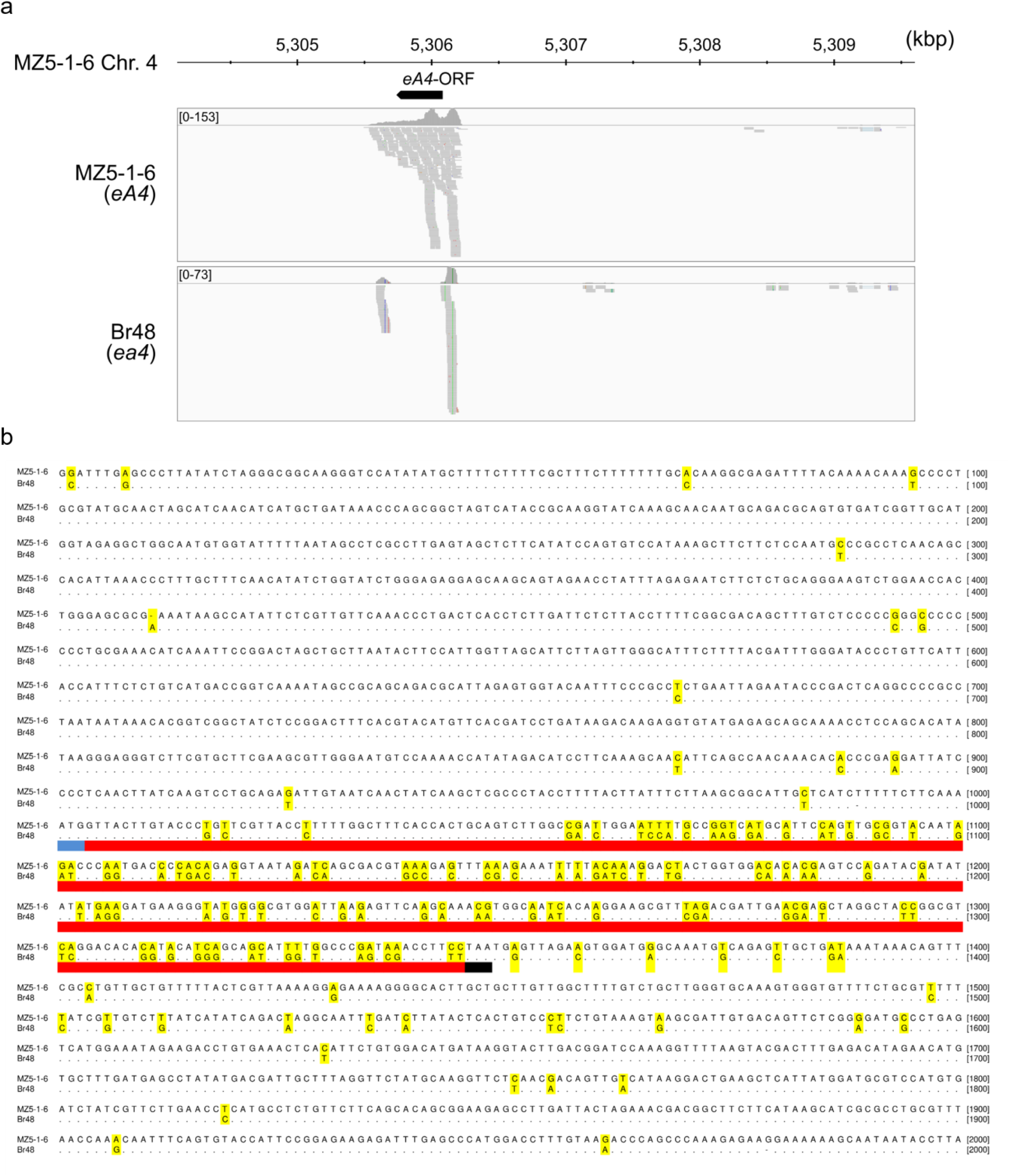
Identification of an *eA4* candidate gene. (a) Alignment of RNA-seq short reads from MZ5-1-6 and Br48 around the *eA4* candidate region in the MZ5-1-6 reference genome. RNA was extracted from primary leaves of barley cv. Nigrate at 24 h after inoculation with each isolate. (b) Nucleotide sequence alignment of *eA4* candidate regions extracted from the whole-genome assemblies of MZ5-1-6 (MoE) and Br48 (MoT). Nucleotides highlighted in yellow indicate sites polymorphic between MZ5-1-6 and Br48. The blue–red–black bar represents the open reading frame (ORF) of the *eA4* candidate gene, with the blue and black segments indicating the start and stop codons, respectively.

**Extended Data Fig. 5.**
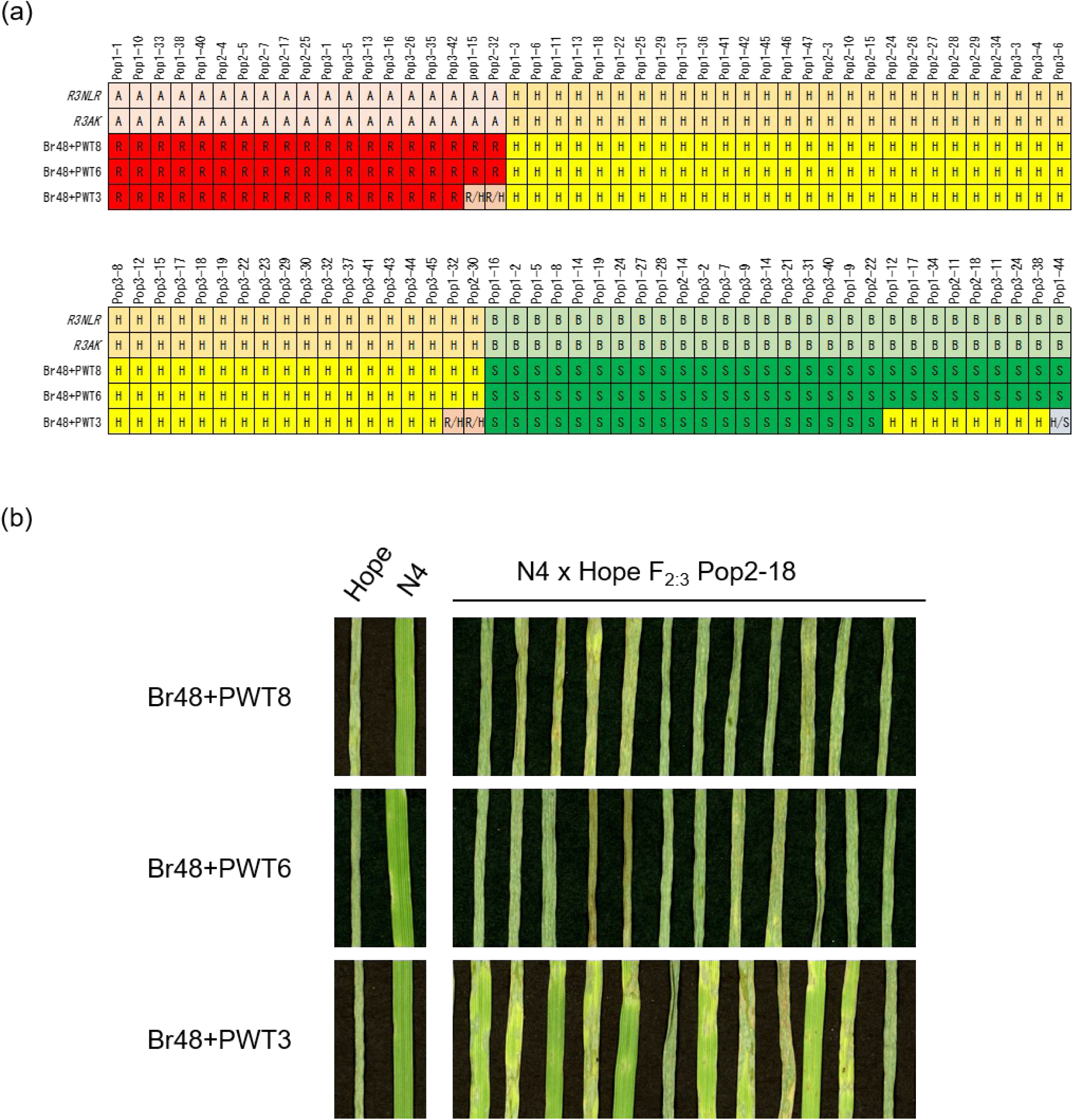
Detection of a resistance gene recognizing *PWT3* alone using the N4 (*Rwt3.6.8*) × Hope (*rwt3.6.8*) population. (a) Segregation of *R3NLR* and *R3AK* (A, present in all; H, segregating; B, absent in all) and reactions to Br48 transformants carrying *PWT8* (Br48+PWT8), *PWT6* (Br48+PWT6), or *PWT3* (Br48+PWT3) (R, all resistant; H, segregating; S, all susceptible) in the F_2:3_ mapping population derived from N4 × Hope. R/H and H/S indicate cases in which the phenotype was difficult to classify as “R or H” and “H or S”, respectively. (b) Responses of individuals of the Pop2-18 F_2:3_ line, which lacks both *R3NLR* and *R3AK*, to the Br48 transformants carrying *PWT8*, *PWT6*, or *PWT3*, five days after inoculation.

**Extended Data Fig. 6.**
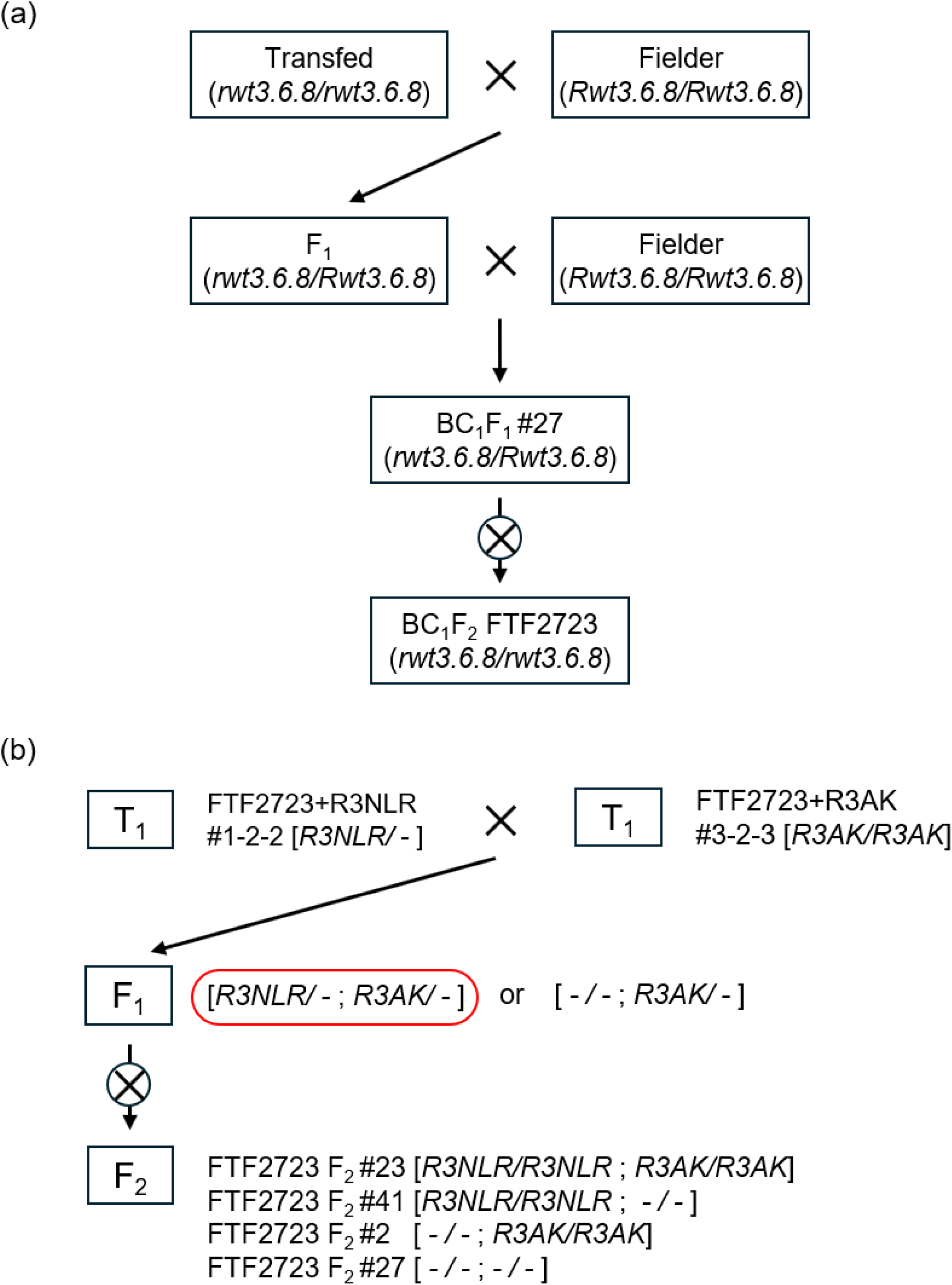
Development of wheat line FTF2723 and its homozygous transformants carrying *R3NLR* alone, *R3AK* alone, both transgenes, and no transgene. (a) Pedigree of FTF2723, a transformable wheat line carrying the *rwt3.6.8* allele. (b) Crossing and selection scheme used to generate homozygous lines.

**Extended Data Table 1.**
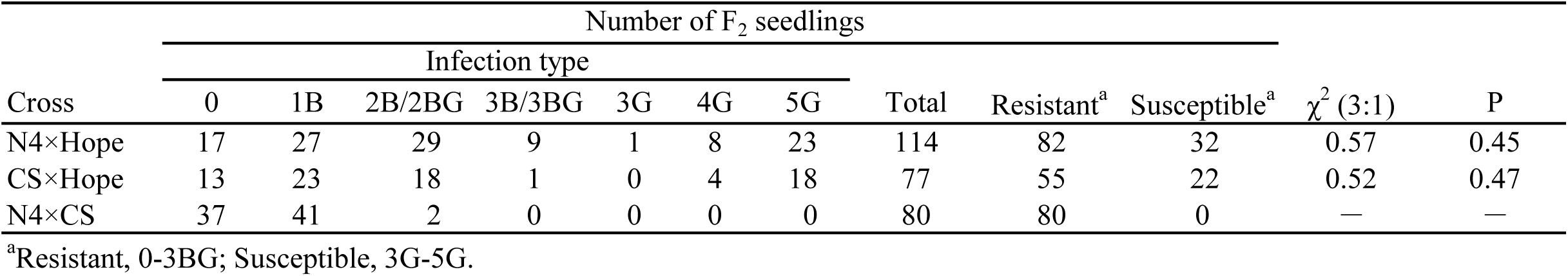
Segregation of reactions to Br48+*PWT8* in F_2_ populations derived from crosses between wheat cultivars.

| Cross | Number of F<sub>2</sub> seedlings | | | | | | | Total | Resistant<sup>a</sup> | Susceptible<sup>a</sup> | $\chi^2$ (3:1) | P |
| --- | --- | --- | --- | --- | --- | --- | --- | --- | --- | --- | --- | --- |
|  | Infection type |  |  |  |  |  |  |  |  |  |  |  |
|  | 0 | 1B | 2B/2BG | 3B/3BG | 3G | 4G | 5G |  |  |  |  |  |
| N4×Hope | 17 | 27 | 29 | 9 | 1 | 8 | 23 | 114 | 82 | 32 | 0.57 | 0.45 |
| CS×Hope | 13 | 23 | 18 | 1 | 0 | 4 | 18 | 77 | 55 | 22 | 0.52 | 0.47 |
| N4×CS | 37 | 41 | 2 | 0 | 0 | 0 | 0 | 80 | 80 | 0 | — | — |

<sup>a</sup>Resistant, 0-3BG; Susceptible, 3G-5G.

