## Supplementary Tables for "A wheat immune receptor pair composed of NLR and MLKL confers stable resistance to pathotypes of the blast fungus by recognizing three effectors"

Supplementary Table S1. *Pyricularia oryzae* isolates used in the present study.

| Isolate | Host | Pathotype | Locality | Year | Accession number of Sequence Read Archive (SRA) |
|---|---|---|---|---|---|
| Ina168 | *Oryza sativa* | MoO | Japan | 1958 | DRR195603 |
| Ken53-33 | *O. sativa* | MoO | Japan | 1953 | DRR413352 |
| PO12-7301-2 | *O. sativa* | MoO | Indonesia | 1973 | DRR413353 |
| PH297 | *O. sativa* | MoO | Philippines | 2012 | DRR413354 |
| 2012-01 | *O. sativa* | MoO | Japan | 1976 | DRR342044 |
| Ao92-06-2 | *O. sativa* | MoO | Japan | 1992 | DRR342045 |
| Ina87T-56A | *O. sativa* | MoO | Japan | 1987 | DRR342046 |
| 85-141 | *O. sativa* | MoO | Japan | 1985 | DRR342047 |
| SL91-48D | *O. sativa* | MoO | Japan | 1991 | DRR342048 |
| 2403-1 | *O. sativa* | MoO | Japan | 1976 | DRR342049 |
| H98-315-1 | *O. sativa* | MoO | Japan | 1998 | DRR342050 |
| 0423-1 | *O. sativa* | MoO | Japan | 1976 | DRR342051 |
| Ina85-182 | *O. sativa* | MoO | Japan | 1985 | DRR342052 |
| Br15 | *O. sativa* | MoO | Brazil | 1990 | DRR893802 |
| VHG4.5 | *O. sativa* | MoO | Vietnam | 1996 | DRR893803 |
| GFSI1-7-2 | *Setaria italica* | MoS | Japan | 1977 | DRR413355 |
| NRSI2-2-2 | *S. italica* | MoS | Japan | 1977 | DRR413356 |
| NRSI3-1-1 | *S. italica* | MoS | Japan | 1977 | DRR413357 |
| NNSI3-2-1 | *S. italica* | MoS | Japan | 1984 | DRR413358 |
| IN77-16-1-1 | *S. italica* | MoS | India | 1977 | DRR413359 |
| IN77-20-1-1 | *S. italica* | MoS | India | 1977 | DRR413360 |
| KANSV1-4-1 | *S. viridis* | MoS | Japan | 1975 | DRR413361 |
| NI913 | *S. viridis* | MoS | Japan | 1974 | DRR413362 |
| NI986 | *Eragrostis lehmanniana* | *Eragrostis* lineage | Japan | 1975 | DRR413371 |
| FSECu1-1-1 | *Er. curvula* | *Eragrostis* lineage | Japan | 1988 | DRR413372 |
| SZECu1-1-1 | *Er. curvula* | *Eragrostis* lineage | Japan | 1988 | DRR413373 |
| SZEC1-1-1 | *Eleusine coracana* | MoE (EC-I) | Japan | 1978 | DRR413363 |
| GFEC1-5-1 | *El. coracana* | MoE (EC-I) | Japan | 1977 | DRR413364 |
| UG77-7-1-1 | *El. indica* | MoE (EC-I) | Uganda | 1977 | DRR413365 |
| UG77-15-1-1 | *El. coracana* | MoE (EC-I) | Uganda | 1977 | DRR413366 |
| UG77-17-1-1 | *El. coracana* | MoE (EC-I) | Uganda | 1977 | DRR893804 |
| Z2-1 | *El. coracana* | MoE (EC-II) | Japan | 1977 | DRR059888 |
| MZ5-1-6 | *El. coracana* | MoE (EC-II) | Japan | 1976 | DRR079313 |
| NP10-17-4-1-3 | *El. coracana* | MoE (EC-II) | Nepal | 1975 | DRR413367 |
| NI1006 | *El. africana* | MoE (EC-II) | Japan | 1975 | DRR413368 |
| NI1011 | *El. boranensis* | MoE (EC-II) | Japan | 1975 | DRR413369 |
| IN77-36-1-1 | *El. indica* | MoE (EC-II) | India | 1977 | DRR413370 |
| IN77-31-1-1 | *El. coracana* | MoE (EC-II) | India | 1977 | DRR1081607 |
| IN77-39-1-2 | *El. coracana* | MoE (EC-II) | India | 1977 | DRR1081608 |
| Br35 | *Urochloa plantaginea* | *Brachiaria* lineage | Brazil | 1990 | DRR079314 |
| TP1 | *Lolium perenne* | MoL | Japan | 1997 | DRR413374 |
| TP2 | *L. perenne* | MoL | Japan | 1997 | DRR079312 |
| TP3 | *L. perenne* | MoL | Japan | 1997 | DRR893805 |
| AK1 | *L. perenne* | MoL | Japan | 1998 | DRR413375 |
| AK2 | *L. perenne* | MoL | Japan | 1998 | DRR893806 |
| LW1 | *L. perenne* | MoL | Japan | 1999 | DRR893807 |
| LW3 | *L. perenne* | MoL | Japan | 1999 | DRR413376 |

| | | | | | |
|---|---|---|---|---|---|
| FI5 | *L. perenne* | MoL | Japan | 1998 | DRR413377 |
| Br58 | *Avena sativa* | MoA | Brazil | 1990 | DRR059887 |
| Br2 | *Triticum aestivum* | MoT | Brazil | 1990 | DRR413378 |
| Br3 | *T. aestivum* | MoT | Brazil | 1990 | DRR413379 |
| Br5 | *T. aestivum* | MoT | Brazil | 1990 | DRR413380 |
| Br8 | *T. aestivum* | MoT | Brazil | 1990 | DRR413381 |
| Br46 | *T. aestivum* | MoT | Brazil | 1990 | DRR413382 |
| Br48 | *T. aestivum* | MoT | Brazil | 1990 | DRR893808 |
| Br49 | *T. aestivum* | MoT | Brazil | 1990 | DRR413383 |
| Br50 | *T. aestivum* | MoT | Brazil | 1990 | DRR413384 |
| Br108.1 | *T. aestivum* | MoT | Brazil | 1992 | DRR413385 |
| Br115.7 | *T. aestivum* | MoT | Brazil | 1992 | DRR413386 |
| Br115.12 | *T. aestivum* | MoT | Brazil | 1992 | DRR413387 |
| Br116.5 | *T. aestivum* | MoT | Brazil | 1992 | DRR079310 |
| Br118.2 | *T. aestivum* | MoT | Brazil | 1992 | DRR079311 |
| Br126.1 | *T. aestivum* | MoT | Brazil | 1992 | DRR413388 |
| Br127.1 | *T. aestivum* | MoT | Brazil | 1992 | DRR413389 |
| Br127.11 | *T. aestivum* | MoT | Brazil | 1992 | DRR413390 |
| Br130.8 | *T. aestivum* | MoT | Brazil | 1992 | DRR413391 |
| Br130.9 | *T. aestivum* | MoT | Brazil | 1992 | DRR413392 |
| BTMP-2(b) | *T. aestivum* | MoT | Bangladesh | 2017 | DRR893809 |
| BTMB-5(b) | *T. aestivum* | MoT | Bangladesh | 2017 | DRR893810 |
| BTGP-6(e) | *T. aestivum* | MoT | Bangladesh | 2017 | DRR893811 |

Supplementary Table S2. Primers used in the present study.

| Objectives | Primer name | Sequence (5' -> 3') | Description |
|---|---|---|---|
| Mapping and cloning of *PWT8* | HK1_F | CTTGCCATGCTTGGAGACTT | Presence/absence marker (present in MZ5-1-6) |
| | HK1_R | ACTGGGGTTTCGCACTGACT | |
| | HK2_F | TCCTTTTGTTCAGATCAACACACG | Presence/absence marker (present in MZ5-1-6) |
| | HK2_R | GAATGCTGCCACACGGAAG | |
| | HK3_F | CGTCAAAAATGATTTCGGTC | Presence/absence marker (present in MZ5-1-6) |
| | HK3_R | TTTACATGCAACAAGACCCAG | |
| | HK4_F | GTAGATGGGTATGCAGCATGGT | Presence/absence marker (present in MZ5-1-6) |
| | HK4_R | CAAAGCCCCAACTTTTGAGG | |
| | HK5_F | GGTAGGTATCCGCCCAAATA | Presence/absence marker (present in MZ5-1-6) |
| | HK5_R | CACACTACAATTCGCCCAAA | |
| | HK6_F | AGTAGGCCTCTGTGCTACGG | Presence/absence marker (present in MZ5-1-6) |
| | HK6_R | GCACTGCTGTCGGTTCATAA | |
| | HK7_F | TTCATTGTCTTATGTACACCGAG | Presence/absence marker (present in MZ5-1-6) |
| | HK7_R | ATTCGGTAGTTGGTTTGCTG | |
| | HK8_F | CACGTAACTCACCTGGCTCA | Presence/absence marker (present in MZ5-1-6) |
| | HK8_R | ATTGCTGCGATCGGTAAAAA | |
| | HK9_F | TCCGGAATCATGGTCCTAAA | Presence/absence marker (present in MZ5-1-6) |
| | HK9_R | AATCCTATCCCGGCTGTACC | |
| | HK10_F | GAACAAGGGCACCATCAAG | Presence/absence marker (present in MZ5-1-6) |
| | HK10_R | GTGGGCGATGTTGTCAAAG | |
| | HK11_F | GATAGCGGACATATGCTAACC | Presence/absence marker (present in MZ5-1-6) |
| | HK11_R | AACCCATGAACTGGTATCAGT | |
| | HK12_F | GTGGAAGATTGATCGAGGGA | Presence/absence marker (present in MZ5-1-6) |
| | HK12_R | TTAAGGTAATCCTTCCGCAA | |
| | HK13_F | GTTATTTTTGGGTGCCGAAC | Presence/absence marker (present in MZ5-1-6) |
| | HK13_R | GCGACGTTGTTTGTCTGAAA | |
| | HK14_F | GGGTCGTCATGATGGCTATT | Presence/absence marker (present in MZ5-1-6) |
| | HK14_R | GCAGCATTCCGACTTAATCC | |
| | HK15_F | CTGTGCAGCACCACAGGA | Presence/absence marker (present in MZ5-1-6) |
| | HK15_R | TGGACGTCTCAAAGTCAGGA | |
| | HK16_F | CGACAGCCAAGTCCCATATT | Presence/absence marker (present in MZ5-1-6) |
| | HK16_R | CATTTTCTCCCACTTGCACA | |
| | HK17_F | CAAAAGGTCTCCTCGACCAC | Presence/absence marker (present in MZ5-1-6) |
| | HK17_R | AATGCAGCTTCACGACAAAC | |
| | HK18_F | GGCTCGGTACATTCAAGGAC | Presence/absence marker (present in MZ5-1-6) |
| | HK18_R | TAATCCTGATCCTCTTGCTCATAT | |

| | | | |
|---|---|---|---|
| | FP | GCTTATGGATCCCTGCGCATC | for fosmid-end sequencing |
| | RP | ATGGACGGTTCCGTGTTTTGG | |
| | PWT8_F | TCTCGTTGTTCAAACCCTGA | for amplification of a 1,047-bp fragment containing the *PWT8* ORF |
| | PWT8_R | CAAGCAGACAAAAGCCAACA | |
| | PWT8-ORF_F | CGGCATTGCTCATCTTTTTC | Presence/absence marker (present in MZ5-1-6) |
| | PWT8-ORF_R | ACTCTGACATTTGCCCATCC | |
| | PWT8-PA_F | ATCAAGCTCGCCCTACCTTT | Presence/absence marker (present in MZ5-1-6) |
| | PWT8-PA_R | AGGTTTATCGGGCCAAAATG | |
| | PWT8-BamHI-Br48-DraI-MZ_R | CAAGCAGACAAAAGCCAACA | CAPS marker used in combination with the PA-F primer |
| Cloning of *Rwt8* | wmc432_F | ATGACACCAGATCTAGCAC | SSR marker[a] |
| | wmc432_R | AATATTGGCATGATTACACA | |
| | gwm337_F | CCTCTTCCTCCCTCACTTAGC | SSR marker[a] |
| | gwm337_R | TGCTAACTGGCCTTTGCC | |
| | RT2_F | TGTCTCCACACCCTGTCGTA | CAPS marker (KpnI) |
| | RT2_R | ATAAAAACGTCGCTGCCAAT | |
| | RT5_F | CTGATCTGGAGGTGCTCAGG | CAPS marker (Hpy99I) |
| | RT5_R | GACTTTCCACCATGGCGATT | |
| | RT8_F | TGTTGCGGCACAGATACTTG | CAPS marker (MspI) |
| | RT8_R | TGCTTGGTTTGATCATTGGA | |
| | YU18_F | ATACAAAATGCGTGTGTAGAGTTCA | Amplification Refractory Mutation System (ARMS) marker |
| | YU18-CS_R | TGTTCCATCACGAGACGAAG | (present in CS and N4, absent in Hope) |
| | YU18-Hope_R | TGTTCCATCACGAGATGAAG | (present in Hope, absent in CS and N4) |
| | YU47_F | GCTCAAAAAACTGGACAGAAGG | CAPS marker (EcoRV) |
| | YU47_R | GTAGTGTGCAACACGCATGG | |
| | NLR17_F | GCTACATCGATTGTTTAGAGGT | Presence/absence marker (present in CS and N4) |
| | NLR17_R | CAGCAAAAAAGACACTTTCC | |
| Protoplast assay | InF-PWT8dSP_F | TGTGTGTGCAGATCGATGGATTGGAATTTTGCCGG | for In-Fusion cloning of *PWT8* without its signal peptide |
| | InF-PWT8dSP_R | GGAAATTCGAGCTCGTTAGGAAGGTTTATCGGGC | |
| | InF-R3NLR_F | TGTGTGTGCAGATCGATGGCGGCGGCTCTTGGT | for In-Fusion cloning of *R3NLR* |
| | InF-R3NLR_R | GGAAATTCGAGCTCGTTACTCGTATCGTACAGATAA | |
| | InF-R3AK_F | TGTGTGTGCAGATCGATGGATCTAGTGGGCAGC | for In-Fusion cloning of *R3AK* |
| | InF-R3AK_R | GGAAATTCGAGCTCGTTAGCCTTTCTTCTTAAACGA | |
| Wheat transformation | hpt_F | GTGTCACGTTGCAAGACCTG | for detection of the *HPT* transgene[b] |
| | hpt_R | GATGTTGGCGACCTCGTATT | |

| | | | |
|---|---|---|---|
| | R3NLR-CK_F | GAAAGCATGGAATGGGAAAA | for detection of the *R3NLR* transgene |
| | R3NLR-CK_R | CCTAAGAAAACCACGGTCCA | |
| | R3AK-CK_F | CTCAACGACCTGGAGGAGAC | for detection of the *R3AK* transgene |
| | R3AK-CK_R | CACCGCGGGATCTCTAATAA | |
| Distribution analysis of *R3NLR* and *R3AK* | R3NLR-PA_F | GTTCCCCCAGCTAAAACTCC | for amplification of a partial sequence of *R3NLR* in *T. aestivum* accessions from genomic DNA |
| | R3NLR-PA_R | GCATGTTGGAGCAGAGCATA | |
| | R3AK-PA_F | CCTCTTGGGTAAGCTGGATG | for amplification of a partial sequence of *R3AK* in *T. aestivum* accessions from genomic DNA |
| | R3AK-PA_R | GGAACCAGCCCCAGTATTGT | |

[a]Somers, D. J., Isaac, P. & Edwards, K. A high-density microsatellite consensus map for bread wheat (*Triticum aestivum* L.). *Theor. Appl. Genet.* **109**, 1105–1114 (2004).

[b]Abe, F. et al. Genome-edited triple-recessive mutation alters seed dormancy in wheat. *Cell Rep.* **28**, 1362–1369.e4 (2019).
